# Imagery priming of binocular rivalry is not a reliable metric of individual differences in the subjective vividness of visualisations

**DOI:** 10.1101/2024.10.28.620764

**Authors:** Loren N. Bouyer, Dietrich S. Schwarzkopf, Blake W. Saurels, Derek H. Arnold

## Abstract

Most people can imagine images that they experience within their mind’s eye. However, there are marked individual differences, with some people reporting that they are unable to construct mental images at all (aphantasics), and others who report having imagined experiences that are as realistic as seeing (hyper-phantasics). The vividness of imagined images is most often measured via subjective self-report. Chang and Pearson (2018), however, have suggested that a binocular rivalry (BR) protocol can be used as an *objective* measure. They found that pre-imagining a moving input could enhance performance on an *objective* probe detection task when probes were embedded in imagery consistent inputs, as opposed to imagery inconsistent inputs. To date, nobody has assessed if this type of *objective* imagery priming can be used to predict the vividness of different people’s visualisations. Here, we report that imagery priming of *objective* sensitivity to probes within static BR inputs *does not* correlate with the typical ratings people use to describe the vividness of their visualisations (a between participants effect). However, *objective* priming of sensitivity to probes embedded in BR inputs was greater on trials when participants reported that their pre-imagined experience had been relatively vivid (a within participants effect). Overall, our data suggest that while imagery can prime objective sensitivity to probes during BR, there is currently no strong evidence that this effect can be used as an objective method of predicting the subjective vividness of different people’s visualisations.

**Highlights:**

➢ There are marked individual differences in the subjective intensity of visual imagery
➢ Most measures of the subjective intensity of visual imagery are subjective
➢ A binocular rivalry probe-detection protocol has been suggested as an objective measure
➢ The authors find that there is only a very small predictive relationship between outcomes in a binocular rivalry task and participant’s reported imagery strength
➢ We still lack an objective measure that reliably predicts the subjective intensity of visual imagery

## Introduction

### Visual Imagery

Sensory imagination is the ability to form a sensory experience within the mind without relevant external input (Pearson, 2019). Imagined visual experiences are often referred to as having a ‘mind’s eye’ (Pearson, 2019). For most people visual imagery is described as an integral tool used frequently in daily life. For example, it is used in memory operations, such as when recalling lecture materials (Dijkstra et al., 2022; Schacter et al., 2012). Sensory imagination can also facilitate social learning, prediction dissonance and planning (Barron et al., 2020; Barron et al., 2013; Dijkstra et al., 2022).

Sensory imagination is a popular tool used in psychological interventions. Therapists use visual imagery to reduce stress, and as a form of exposure therapy to treat phobias (Hirsch & Holmes, 2007; Holmes & Mathews, 2010; Reddan et al., 2018).

Sports psychologists use imagination to improve the performance of athletes (Veraksa & Gorovaya, 2011), and counsellors use imagination to prepare for future conflict and social interactions (Grzanka & Moradi, 2021). Therefore, by understanding the function, processes, and individual differences that characterise visual imagery, we can better tailor these types of intervention to different people.

### The Vividness of Visual Imagery

The ability to have imagined visual experiences varies dramatically between people. Some people report being aphantasic; unable to produce any mental images (Zeman et al., 2015; Zeman, 2024). While aphantasia can be acquired after neurological trauma (Zeman et al., 2010), some people report that they have never been able to conjure an imagined visual experience. These people are referred to as having congenital aphantasia (Zeman et al., 2015). Congenital aphantasia is unusual, occurring in less than 5% of the population (Zeman et al., 2015). However, many aphantasics do not realise that their cognitive profile is unusual, potentially at least in part because they have formed ways to compensate for their lack of visual imagery.

Another segment of the population, by contrast, report being hyperphantasic; able to produce imagined visual experiences that seem as real to them as actual seeing (Galton, 1880a; Zeman, 2024). These experiences can be voluntary or involuntary. Extreme instances of involuntary realistic experiences can be seen in synaesthetes (Arnold et al., 2012; Galton, 1880b; Simner etal., 2006) and in people with post-traumatic stress disorder (e.g. Morina et al., 2013). Between the two extremes of aphantasia and hyperphantasia, there seems to be a spectrum of imagination vividness (Dijkstra et al., 2022). These individual differences in the subjective vividness of imagined visual sensations seems to be related to differences in brain morphology and neural activity (Arnold etal., 2024; Bergmann et al., 2016; Dijkstra et al., 2017; Zeman, 2024).

#### Current Measures of the Vividness of Visualisations

A substantive issue for this area of research is how to best measure the subjective vividness of imagined visual experiences. The current standard measure is a self-report questionnaire; the Vividness of Visual Imagery Questionnaire 2 (VVIQ2) (Marks, 1995). However, this measure is entirely subjective (Zeman, 2024).

Moreover, there is no standardisation of ratings between people; so one person’s low ratings might coincide with another person’s high ratings despite both people having matched imagined experiences. This suggests that if we rely on subjective metrics like the VVIQ2 there will likely be some measurement error due to ambiguity in relation to how different people interpret the available choice options.

The experiences of the first author of this paper, highlight concerns regarding the ambiguity of the VVIQ2 (Marks, 1995). She is a self-described congenital aphant, who feels that she has never had an imagined visual experience during her waking life. However, when she first completed the VVIQ2 (Marks, 1995) her responses were stereotypical. She had interpreted the idea of seeing a mental image as a literary expression that related to the effort expended in trying to recollect facts, rather than as questions about how vividly she had experienced an actual mental image.

The VVIQ2 was never intended to be a metric for the diagnosis of Aphantasia, so there is growing awareness that a better objective instrument is desirable (e.g. Blomkvist & Marks, 2023; Schwarzkopf, 2023). Blomkvist and Marks (2023) have pointed out that the arbitrary and diverse criteria (in terms of VVIQ2 scores) used to categorise people as aphantasic in different studies could compromise cross-study comparisons and research into the underlying causes of aphantasia. They also suggest that self-diagnoses from online versions of the questionnaire could be harmful to the welfare of people mistakenly identified as aphantasic (Blomkvist & Marks, 2023). Therefore, it is important to find and validate a more reliable measure of the subjective intensity of imagined visual experiences than the VVIQ2 (Marks, 1995), and ideally an objective task that predicts this.

Some studies have identified candidate tasks that might predict the subjective intensity of imagined visual experiences (e.g. Chang & Pearson, 2018; Cui et al., 2007; Kay et al., 2022). Of these, perhaps the leading candidate is a binocular rivalry (BR) protocol (Chang & Pearson, 2017).

BR can occur when discrepant images project to corresponding points on the retina in each eye (Arnold, 2011; Hemlholtz, 1962; Levelt, 1968). After a brief initial period, one of the two rivalrous images can become suppressed from conscious awareness, with the other image dominating perception. If rivalrous presentations are protracted, the status of the two images can stochastically switch, between being dominant and being suppressed from awareness.

Pre-imagining a rivalrous input can prime participants to report that the pre-imagined input was the first to dominate perception during a subsequent rivalrous presentation (Chang et al., 2013; Pearson et al. 2008). Moreover, this priming effect has scaled with the reported intensity of pre-imagined experiences. This has been true of both the different ratings people use to describe the subjective intensity of their imagined experiences on different trials (i.e., a within subjects effect, e.g., Pearson et al., 2011) and of the average ratings different people have used to describe their typical imagined visual experiences (i.e., a between subjects effect, see Pearson et al., 2011). This last type of finding is particularly intriguing, as it suggests BR priming could be used as a metric to assess an individual’s capacity to have imagined visual experiences. Some studies have, however, failed to detect a between subjects imagery vividness priming effect on BR (see Bergmann et al., 2016, Supplemental Figure S1c; Dijkstra et al., 2019). Moreover, there is an important caveat to consider in relation to this type of evidence. Having people report on perceptual dominance during BR is an entirely *subjective* measure. Without further evidence, imagery priming of BR dominance is just an alternate subjective measure, relative to simply having people report on the subjective vividness of their imagined experiences.

### Objective Tasks, Subjective Tasks and Response Biases

Objective tasks test sensitivity to a ground truth. An experimenter might, for instance, present a test probe above or below fixation and ask a participant to indicate where that input had been presented. As there is an objectively correct answer, the proportion of correct answers should have a floor that approximates chance task performance (Green & Swets, 1966; Yarrow et al., 2011). If a person has no sensitivity to test probes they should be reduced to guessing as to their locations, and should guess correctly on ∼50% of trials (if there is 1 incorrect response option, see Green & Swets, 1966).

When a participant has no sensitivity they might have a systematic response bias (Green & Swets, 1966; Yarrow et al., 2011). For instance, if they cannot detect a probe they might always guess that the probe had been presented above fixation. In an objective task, where the experimenter has randomised the test probe location, this bias will not benefit or harm the overall proportion of correct guesses – the participant should still guess correctly at chance on ∼50% of trials.

Researchers are often interested in people’s *subjective* experiences. BR experiments can be used to study which of two rivalrous inputs a participant is experiencing. Though they are being exposed to both, and each is exciting activity in their visual cortex (Brown & Norcia, 1997; Haynes & Rees, 2005), they might only be subjectively aware of one input. Researchers are very interested in the neural processes that govern these *subjective* experiences. A key characteristic is that, because there is no *objectively* correct answer, the percentage of trials on which a participant reports having a particular type of experience (such as seeing a pre-imagined rivalrous input, e.g. Andermane et al., 2020; Chang et al., 2013; Dijkstra et al., 2019; Keogh & Pearson, 2011, 2014, 2017, 2018; Keogh et al., 2020; Maróthi & Kéri, 2018; Pace et al., 2023; Rademaker & Pearson, 2012; Shine et al., 2015) can reasonably range between 0 and 100%.

In studies of subjective experience, researchers quantify response biases – any tendency to report one type of experience more often than another. A key problem is that it can be difficult to interpret the cause(s) of a response bias (Gallagher et al., 2019; Storrs, 2015; Yarrow et al., 2011). People might, for instance, be biased to report seeing a pre-imagined input during rivalry because visualization has primed the visual system to be more responsive to that input. Alternatively, the participant might be unsure which input they are seeing, but be more willing to reporting ‘seeing’ a pre-imagined input – a decision level bias as opposed to a perceptual effect. As both scenarios can result in matched patterns of behavioural response, in the absence of further evidence these effects cannot confidently be said to have a perceptual origin (Gallagher et al., 2019; Storrs, 2015; Yarrow et al., 2011).

This type of criticism of BR imagery research has been countered by including non-rivalrous catch trials, with stimuli in a number of studies consisting of a common spatial mix of both inputs (i.e. some regions of the input could be left titled, and other regions right tilted). These stimuli are shown to both eyes, so BR is not experienced (e.g. Chang et al., 2013; Keogh & Pearson, 2014; 2017; Pearson et al., 2008, 2011). The idea is that if participants are not biased to report seeing a pre-imagined input on these ‘catch’ trials, then the bias observed on rivalrous trials can safely be ascribed to perception.

However, there are important differences between the perceptual experience of rivalrous and non-rivalrous trials. First, rivalrous presentations involve a sense of lustre at the site of rivalry that is absent from non-rivalrous displays (Wendt & Faul, 2022). Second, there is ambiguity in relation to when an input has become dominant during rivalry, which requires a participant to reference their experiences against a subjective criterion. Rivalrous presentations typically involve an initial period when either both images can be seen as a transparent display, or during which parts of both images can be seen (Anderson, et al., 1978). Dominance then tends to develop gradually, often in a non-linear manner. One input can, for instance, initially seem to fade, only to reassert itself in perception and be the first to dominate the other input (Anderson, et al., 1978). So, a participant needs to index their changing experiences against a subjective criterion to decide when an input has completely disappeared.

These decisions are not typically needed on catch trials (e.g. Change et al., 2013; Keogh & Pearson, 2014; 2017; Pearson et al., 2008, 2011) as perception should both clearly consist of parts of both images, and it should be unchanging.

Researchers have also attempted to counter the idea that BR protocols might serve as an alternate *subjective* measure of the vividness of imagery by examining the specificity of imagery-rivalry interactions (e.g. Bergmann et al., 2016; Pearson et al., 2008). The first thing to note about these manipulations is that they are still a measure of a subjective decision process. In one example, in a small group, Pearson et al. (2008) found that pre-imagining an oriented Gabor had a similar impact as seeing an oriented Gabor before a rivalrous presentation. Both effects were tuned for similarity in terms of the orientations of rivalrous inputs and pre-imagined primes. By contrast, Bergmann et al. (2016) only found evidence for a coarser orientation tuning of imagery priming in a subset (∼1/3) of a much larger group of participants (see their Figure 3, and compare to Pearson et al., 2008 Figure 5). These data are interesting, but there is no reason a cognitive bias of a subjective experience (e.g. Roseboom & Arnold, 2011) should not have a similar reliance on similarity relative to a perceptual effect.

In relation to location, Pearson et al. (2008) found that pre-imagining an oriented Gabor had a similar impact as paying attention to a physical oriented Gabor. They both primed participants to report dominance of the primed rivalrous input when rivalrous displays were presented in the same location, but not otherwise. This contrasts with Bergmann et al. (2016, see Supplementary Results) who did not find any evidence for greater imagery priming when people were asked to pre-imagine inputs in the same as opposed to a different location (F_1,30_ = 1.36, p = 0.25). It is not clear if participants in either study were asked if they felt they could imagine inputs as being externally localised within the world, as opposed to being located entirely within their mind’s eye. This is a concern as only a subset of people may be able to imagine inputs as being externally located (Salfaro et al., 2024; Schwarzkopf, 2024). Regardless, overall these data provide weak evidence for a tight spatial dependency of imagery priming, and there is no reason why a cognitive bias could not be specific to an attended or to an imagined location in any event.

### Imagery priming of an objective test of visual sensitivity

Mindful of criticism concerning the subjectivity of imagery-rivalry interactions, Chang and Pearson (2017) extended the visual imagery BR protocol by adding a probe detection task. As described above, this *can* deliver an objective measure of visual sensitivity. A probe can be added above or below fixation within one of the two rivalrous inputs, and participants must indicate in which position the probe was presented. The important finding is that participants are more likely to correctly identify a probe’s location when it is embedded in an input that is currently dominating perception (Alais & Parker, 2006; Wales & Fox, 1970). Moreover, this objective priming effect wanes as a transition of perceptual dominance nears and suppression weakens (Alais et al., 2010).

Chang & Pearson (2017) found that people were overall more sensitive to probes embedded in pre-imagined inputs. The authors argued that this had served as an objective index of imagery (Chang & Pearson, 2017). In the original study, however, this effect was not examined on a between participants basis – to assess if it could predict individual differences in the subjective vividness of imagined visual sensations. Another point of ambiguity was caused by the study protocol. Probes were presented throughout rivalrous presentations, and no dominance reports were collected. These data could therefore have been driven by imagery biasing participants to selectively attend to one of the two rivalrous inputs, with more probes being detected within attended inputs before the onset of rivalry.

### Our Goal: To identify a reliable objective metric of individual differences in the subjective vividness of visualisations

Our overarching goal is to identify an objective metric that can be used to reliably detect individual differences in the subjective vividness of imagined visual sensations. The leading candidate task is a BR protocol, where imagery may prime sensitivity to probes embedded in rivalrous presentations (Chang & Pearson, 2017). We chose to investigate this issue using a protocol where participants report on the onset of dominance - so we would know if probes were presented to the reportedly dominant or to the suppressed eye.

### Methods Participants

A preliminary study conducted in our lab had suggested an r of ∼0.35 between individual VVIQ2 scores and the power of correlated measures of oscillatory brain activity (Arnold et al., 2023). Using this as a guide as to what relationship we might hope to detect between individual VVIQ2 scores and imagery priming of probe detections embedded in inputs undergoing BR, we conducted a power analysis based on an α = 0.05 and β = 0.8, and this recommended a sample size of 62 to measure our target correlation. To allow for extraneous issues such as equipment failure, participant dropout etc., we preregistered a total of 80 participants (https://aspredicted.org/uv2wb.pdf). When asked for their gender identity the overall sample consisted of 21 men, 57 women, and 2 others, with ages ranging from 17 to 52 (*M*=22, *SD*=7). These consisted of 50 volunteers recruited via an undergraduate student research participation scheme, where students received course credit in return for their participation, 16 participants recruited through word-of-mouth, and 14 participants who were compensated with $40 for their time. All participants had normal or corrected-to-normal visual acuity (i.e., they wore glasses or contact lenses if they needed them to read).

Of the participants, three women did not complete the BR task due to technical issues or an inability to detect stimuli. The demographics of the final sample therefore changed, to 21 men, 54 women, and 2 others, with a mean age of 22 (*SD*=7).

All data and the analysis script that has informed the results described in this paper are publicly available.

### Materials Apparatus

Testing took place in a darkened room. A chinrest was used to ensure a constant viewing distance of 50 cm from the monitor. Participants wore anaglyph glasses, with a green filter covering the left eye and a red filter covering the right eye. This results in wavelengths of light associated with the colour red (CIE: x=0.61, y =0.38) being filtered (appearing to be black) when viewed through the red filtered right eye, and white when viewed through the green filtered left eye, and vice versa for wavelengths of light associated with the colour blue (x=0.14 y=0.64). The red and blue colours were selected as the colours most effectively filtered when viewed through the red and green anaglyph glasses – as measured by a CRS ColorCal colorimeter. This ensured that there was no cross talk. Visual stimuli and task instructions were presented on an ASUS VG248QE 3D Monitor (1920 x 1080 pixels, refresh rate: 60 Hz) driven by a Cambridge Research Systems ViSaGe stimulus generator and custom MATLAB R2015b software.

### Questionnaire

Prior to the formal experiment, participants completed the Vividness of Visual Imagery Questionnaire v2 (VVIQ2); a psychometric measure of the vividness of imagined visual experiences (Marks, 1995). This questionnaire consists of 32 items (e.g., *In answering item 8, think of the item mentioned in the following question and rate the vividness of your imagination: a rainbow appears*) each rated on a 5-point scale (from 1. *No image at all. I only know that I was thinking of images*, to 5. *Perfectly realistic, as vivid as real seeing*). Questionnaire scores can therefore range from a minimum of 32 to a maximum of 160.

### BR Stimuli

For the first 31 participants, the luminance of Gabors was calculated from their on-screen properties. The peak luminance of these Gabors were red (CIE: x=0.61, y =0.38, Y=1.22) and blue (x=0.14 y=0.64 Y=1.22), and they were presented against a mid blue/red background (see Figure 1). However, after noticing that the anaglyph glasses were a greater filter of red, peak on-screen red luminance (red CIE: x=0.61, y =0.38, Y=2.11) was increased for the final 49 participants to equate peak red and blue luminance values as recorded through the anaglyph filters (Y=1.22). A between groups t-test was conducted to ensure that this had not resulted in differences in BR priming from imagery (see Results).

**Figure 1.**
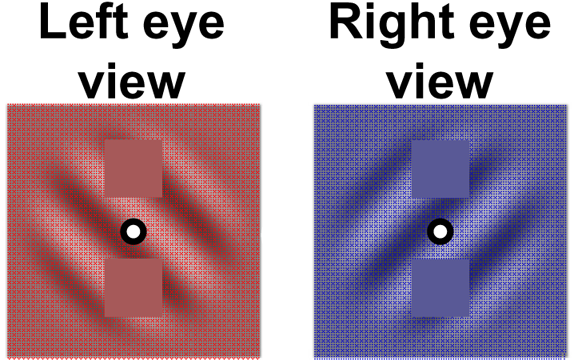
Right-tilted and Left-tilted Gabors, with embedded square regions where probes could be presented.

Red Gabors were orientated 45° left titled from vertical and blue Gabors were orientated 45° right tilted from vertical (see Figure 1). All Gabors had a spatial frequency of one cycle / degree of visual angle (dva), and a spatial constant that subtended 1 dva. The phase of each Gabor was set at random (0-360°) on a trial-by-trial basis, to mitigate against the build-up of retinal after-images.

Rivalry was instigated by alternately presenting Left and Right tilted Gabors respectively to the left and right eyes at the monitor refresh rate. Within the rivalrous Gabors that were presented at the beginning of each trial sequence, there were square regions subtending 1 dva in diameter, centred horizontally within the Gabor and 0.75 dva above and below a central black and white bull’s eye fixation point (that was visible to both eyes). These square regions were set to the mid red or to the mid blue colour of the relevant Gabor. Catch trial Gabors occurred when only one of the two Gabors was presented on the screen. Probe presentations were triggered when participants reported that one of the two rivalrous Gabors had begun to dominate perception. Probes then consisted of the elimination of one of the two mid red or mid blue squares from within a rivalrous Gabor, to be replaced by the completion of the Gabor waveform across that square region (see Figure 2).

**Figure 2.**
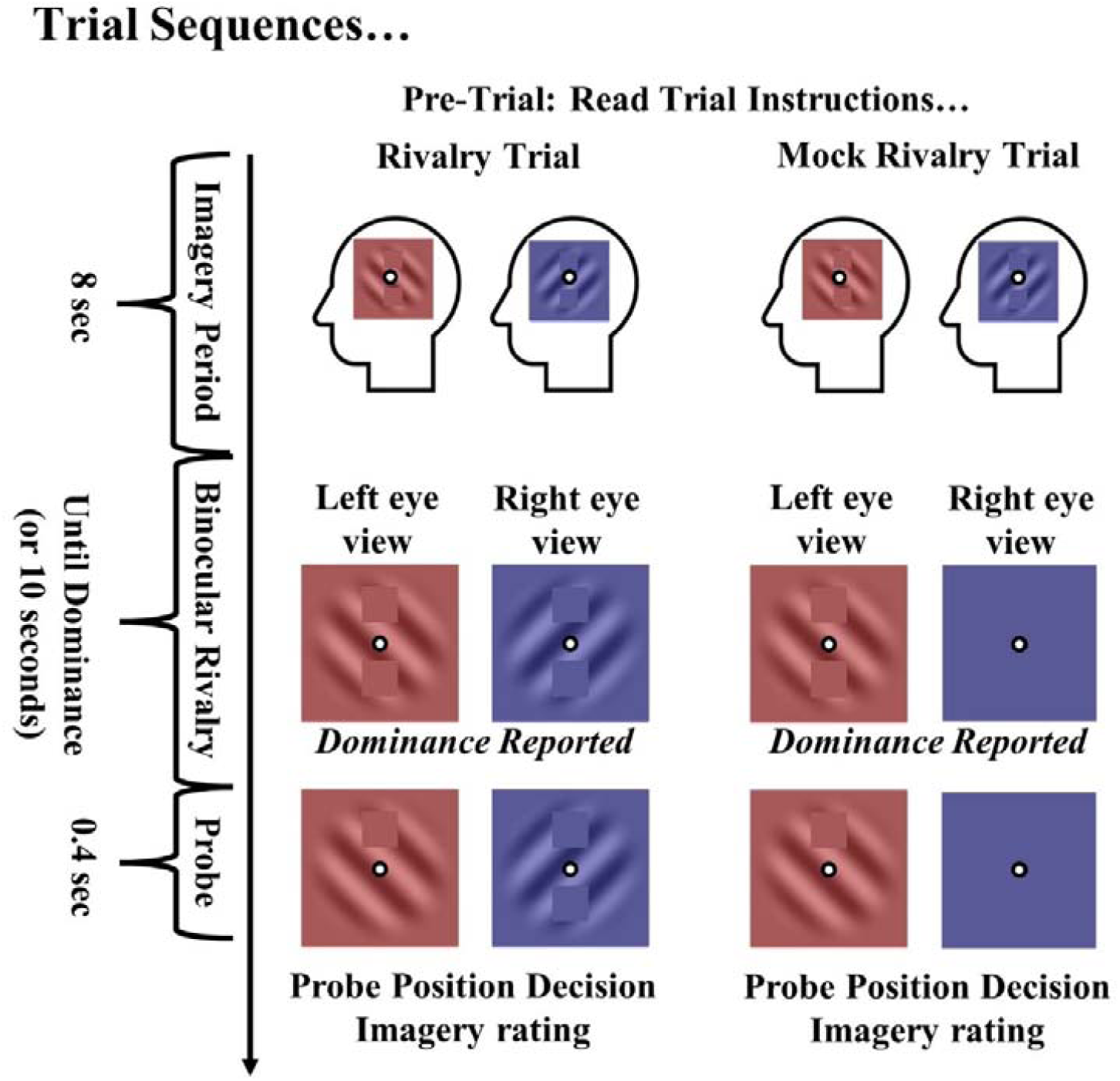
Graphic depicting a Rivalry (left) and a Mock rivalry trial sequence (right). Trials began with an 8 second visual imagery period, during which participants were asked to imagine seeing one of two rivalrous Gabors in as much detail as they could. After the imagination period, either a rivalrous or a Mock rivalrous stimulus presentation began. This persisted until the participant indicated that one of the two rivalrous Gabors had begun to dominate perception, or until 10 seconds had passed. In the former case, a probe would then be presented for 0.4 seconds. In rivalrous presentations this was shown to either eye, whereas in Mock rivalry trials it was always shown within the single Gabor that had been presented. After probe presentations participants would sequentially indicate **1)** where the probe had been presented (above or below fixation), and **2)** how vivid their experience of the pre-imagined Gabor had been. If perceptual dominance was not reported within 10 seconds, the trial was ended and instructions for the next trial were shown.

**Figure 3.**
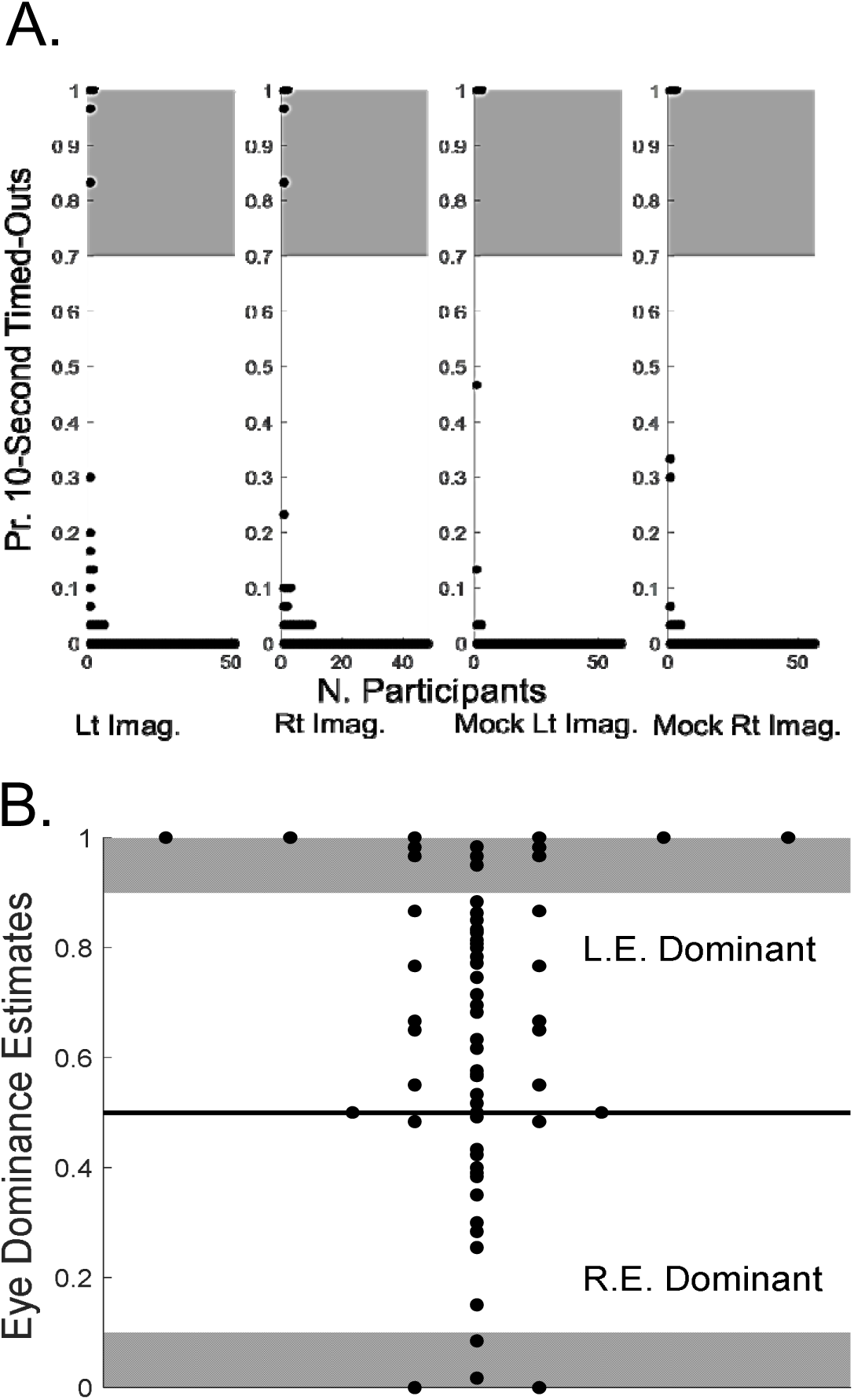
Exclusion of participants based on **A**) time-outs and **B**) eye dominance estimates. **A**) Subplot showing the proportion of 10-second stimulus presentations on which different numbers of people (X-axis) did not report perceptual dominance (Y-axis). The 4 subplots depict data for the four experimental conditions: imagine left or right tilted Gabors before a rivalrous display, and imagine left or right tilted Gabors before a mock rivalry display. The grey shaded area indicates participants who were excluded from analyses on the basis that they did not experience perceptual dominance on a sufficient number of trials. **B.** The proportion of left or right eye dominance reported by different people (Y-axis), averaged across Imagine Left and Imagine Right BR trials. The black horizontal line represents equal reporting of Left and Right eye dominance across all BR trials. Scores above the black horizontal line represent participants who reported more left eye dominance across all BR trials, and scores below the black horizontal line represent participants who reported more right eye dominance across all BR trials. The grey shaded areas represent participants who were excluded from further analyses because they were strongly biased to report perceptual dominance of content in one eye, regardless of Imagined content.

During imagery periods, participants fixated a black and white bull’s eye fixation point (0.6 dva in diameter) centred on a display set to a combination of the mid red and blue Gabor colours (CIE: x=0.18, y =0.09, Y=3.25).

### Procedure

At the beginning of each experimental session, participants completed demographic questions (about age and gender identity) and the VVIQ2. They then completed the BR task.

After adorning their face with anaglyph glasses, participants were familiarised with the appearance and labels of left/right-tilted Gabors (see Figure 1) and of probe stimuli, as viewed through the anaglyph glasses. They then participated in 120 individual trials.

Each trial began with instructions on the screen asking participants to try to imagine in as much detail as they could either a left-tilted or a right-tilted Gabor with the empty squares above and below the fixation point. The fixation point was a central black and white bull’s eye configuration (diameter subtending 0.6 dva, visible to both eyes). They were asked to fixate on this configuration throughout the stimulus presentation. These imagination periods were instigated by the participant clicking a mouse button to begin the trial, and they persisted for 8 seconds. Each type of Gabor was imagined equally often on trials completed in a random order (see Figure 2). After imagination periods, a rivalrous display was presented, centred on the fixation point, or a non-rivalrous catch trial Gabor was presented (equally as often as rivalrous presentations, and in random order). Participants were asked to indicate when they could only see a Left or a Right tilted Gabor. That is, they were asked to indicate the onset of perceptual dominance by clicking the left mouse button for left-tilted Gabor dominance and the right mouse button for right-tilted Gabor dominance. If a participant failed to report the onset of perceptual dominance after 10 seconds, the trial was ended and instructions for the next trial were presented.

Perceptual dominance reports triggered a 0.4 second probe presentation above or below fixation. The probe was presented within the Gabor that was visible to either the Left or Right eye (see Figure 2). The probe was equally likely to be embedded within the pre-imagined Gabor, or within the Gabor that had not been imagined. Participants were then asked to indicate where the probe had been presented (above or below fixation). Participants received feedback on the accuracy of their positional response, and they were then asked to rate the subjective vividness of their pre-imagined Gabor (using the same 5-point scale as used in the VVIQ2, see Marks, 1995).

## Results

### Data Exclusions and Cleaning

As the first 31 participants had been tested with different on-screen peak red luminance values, a between groups t-test was conducted to test for differences in objective probe detections from having pre-imagined Gabors. This refers to the increased proportion of probes that were detected within rivalrous Gabors when they were embedded within pre-imagined Gabors, as opposed to Gabors that had not been pre-imagined. This analysis revealed a non-significant effect (t_55_ = 0.088, p = 0.93) with a Bayes factor analysis (BF_10_ = 0.24) providing moderate evidence for the null hypothesis – that there had been no difference in imagery priming of objective sensitivity to probes between these two groups. The two groups were therefore treated as a single group of participants in all subsequent analyses.

Participants were instructed to report when one of the Gabors had completely dominated perception as quickly as possible by clicking the appropriate mouse button. Those who had not reported on perceptual dominance happening within 10 seconds of the onset of rivalrous input were regarded as having timed out on that trial. On these trials participants either experienced rivalry that was too slow for the experiment to measure, or they had not understood task instructions. Regardless of the cause, timed out trials cannot provide evidence of the impact of imagery. So, participants who timed-out after 10 seconds on 70% or more of rivalrous trials were excluded from further analyses (see Figure 3A, datapoints within grey shaded regions of the two left-side subpanels).

Eye dominance estimates were calculated by determining on what proportion of rivalrous presentations people had reported that the content shown to either their Left or to their Right eyes had first dominated perception, averaged across imagination conditions (see Figure 3B). As these analyses average across the two imagination conditions, they reveal the propensity of a participant to report seeing the content of one eye, independent of any bias from pre-imagining content.

Participants who reported first seeing the content presented to a given eye on 90% or more of all rivalrous trials were excluded from further data analyses, as their experiences of perceptual dominance were evidently determined almost entirely by what eye content had been shown to, as opposed to a competition that could be biased by imagery.

After controlling for time outs and for strong eye dominance, there were 58 participant datasets available for analysis. As a final set of precautions, we excluded datapoints from any analyses that were +-3 standard deviations from any conditional mean, and we only included participant data within a conditional analysis if there were at least five viable trials available from that participant for that condition. The precise numbers of participants informing each analysis are therefore variable, but these can be inferred from the reported degrees of freedom for each test.

After assessing for normality using the Kolmogorov-Smirnov test, we found that the only set of data that violated the assumption of normality was the proportion of imagery consistent dominance (p = 0.008). We therefore used Spearman’s rank correlation when correlating with this variable.

After these exclusions, we confirmed that our participants were not biased to report imagery consistent BR dominance on mock trials (t_52_ = -0.54, p = 0.59, BF_10_ = 0.153).

### Trial-by-trial imagery ratings and VVIQ2 scores were interrelated

To test if trial-by-trial imagery ratings were a relatively reliable measure of the subjective vividness of different people’s imagined visual experiences, we correlated participants’ average trial-by-trial imagery ratings with their VVIQ2 scores. There was a moderate positive relationship (r = 0.408, *p* = 0.001, N = 58, see Figure 4A).

**Figure 4.**
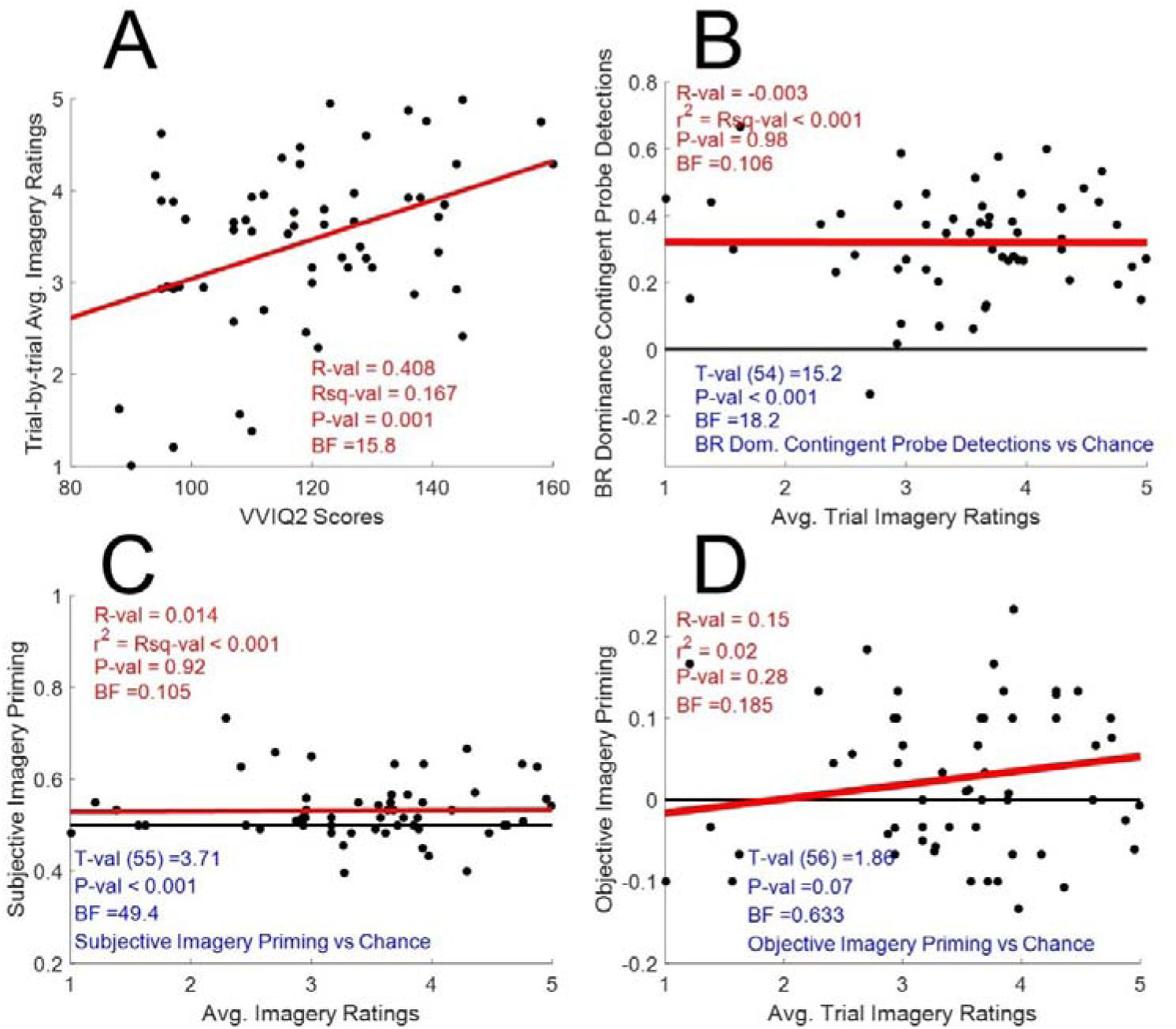
**A)** Scatter plot of individually averaged trial-by-trial imagery ratings (Y-axis) and VVIQ2 scores (X-axis). **B)** Scatter plot of individually averaged trial-by-trial imagery ratings (X-axis) and differences in proportions of detected probes when these were embedded in dominant, as opposed to suppressed inputs during BR. The bold red line depicts a least squares linear trend fit to data and the black horizontal line depicts the difference expected if probe detections were not contingent on BR dominance. **C)** Details are as for Figure 4B, with the exception that individual estimates of subjective imagery priming of BR dominance is plotted on the Y-axis. **D)** Details are as for Figure 4B, with the exception that individual estimates of *objective* imagery priming of probe sensitivity during BR is plotted on the Y-axis.

**Figure 5.**
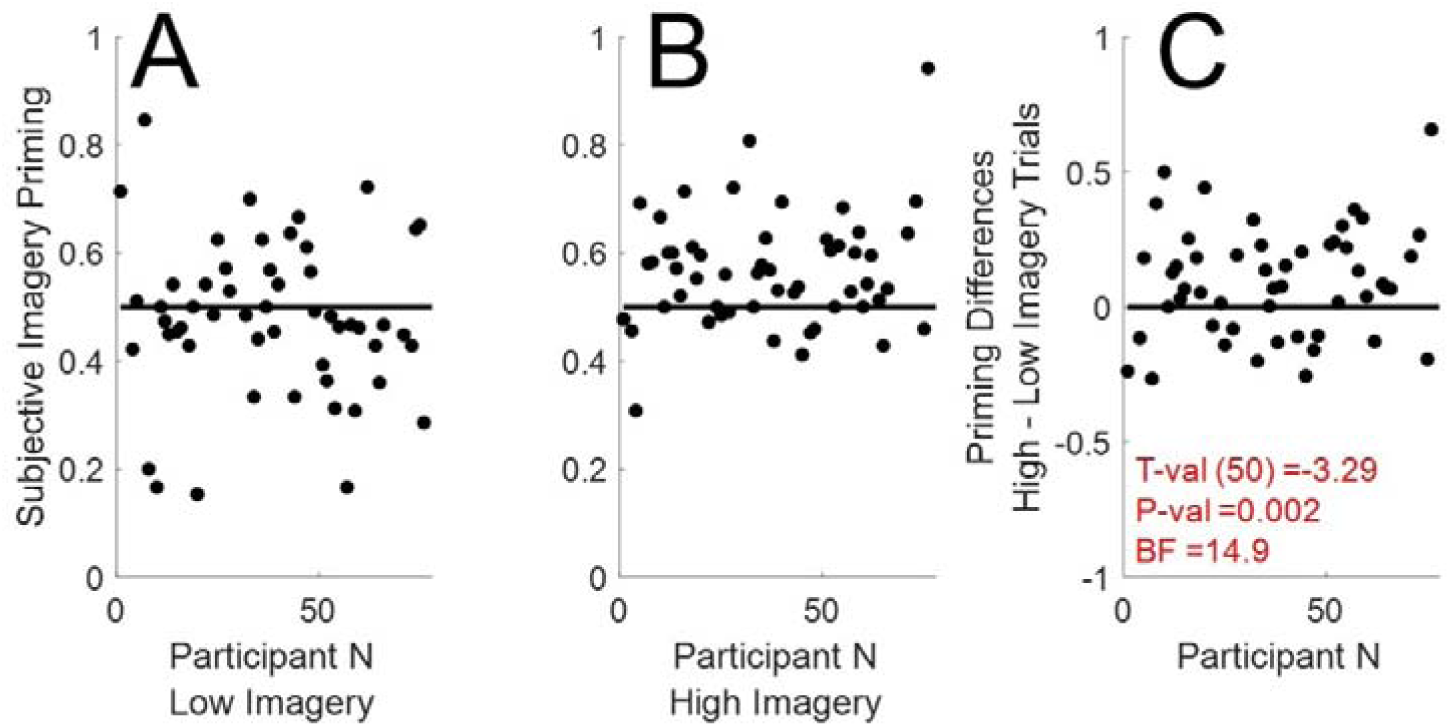
**A.** Scatterplot of subjective imagery priming (the proportion of imagery consistent – the proportion of imagery inconsistent BR dominance reports, Y axis) for different participants (X axis) on low imagery trials. **B.** Details are as for Figure 5A, but for high imagery trials. **C.** Scatterplot of priming differences on high (see Figure 5A) vs low (see Figure 5B) imagery trials (Y-axis) for different participants (X axis).

### BR was instigated, as there was robust evidence of BR and suppression

If participants were experiencing BR, they should detect more probes when these are embedded in the rivalrous Gabor that is reportedly currently dominating perception, as opposed to Gabors that are reportedly currently suppressed from awareness (Alais & Parker, 2006; Wales & Fox, 1970). A paired t-test revealed a robust impact of reported perceptual dominance, such that people detected the positions of more probes embedded in reportedly dominant Gabors (paired t_54_ = 15.2, p < 0.001, BF_10_ = 18.2; see Figure 4B). These data show that these participants were experiencing BR, culminating in the perceptual dominance of one input, and in the perceptual suppression of the other, and that these subjective effects modulated *objective* sensitivity to probe locations.

### Imagery primed subjective dominance during BR

A number of studies have found that pre-imagining content can bias subjective BR dominance reports (e.g. Chang et al., 2013; Pearson et al. 2008). We wanted to assess if this effect was evident in our data. There was an overall impact of pre-imagining a rivalrous input on the probability that it would reportedly first dominate perception (t_55_ = 3.71, p < 0.001, BF_10_ = 49.4; see Figure 4C). So, we have replicated the effect whereby imagery primes *subjective* BR dominance.

### Subjective priming of BR dominance was not correlated with individual differences in vividness of imagery ratings

We found no evidence for a robust correlation between imagery priming of subjective dominance during BR and either the average ratings people used to describe the subjective intensity of their pre-imagined Gabors (r_s_ = 0.014, *p* = 0.92, BF_10_ = 0.105, N = 56; see Figure 4C) or their VVIQ2 scores (r_s_ = 0.017, p = 0.9, BF_10_ = 0.106, N = 56). Note that these data relate to between subjects effects, and they only speak to the impact of imagery on *subjective* dominance during BR.

### Objective priming of sensitivity during BR was not correlated with individual differences in vividness of imagery ratings

We found no robust evidence for a correlation between imagery priming of successful probe position decisions (an objective measure of sensitivity) and either the average ratings different people used to describe the subjective intensity of their pre-imagined Gabors (r = 0.15, *p* = 0.28, BF_10_ = 0.185, N = 57; see Figure 4D) or their VVIQ2 scores (r = 0.039, p = 0.77, BF_10_ = 0.107, N = 57).

#### Imagery did not improve overall objective sensitivity to probes

To assess if pre-imagining Gabors had improved objective sensitivity to probes overall, regardless of imagery vividness ratings, we tested for a difference in the proportions of successful probe position decisions when probes were embedded in pre-imagined Gabors, as opposed to when probes were embedded in Gabors that had not been pre-imagined. We found no evidence for an advantage for probes embedded in pre-imagined Gabors (t_51_ = 1.86, p = 0.07, BF_10_ = 0.633; see Figure 4D).

### There were robust within-participant imagery priming effects, of subjective dominance and objective sensitivity

Imagery had an impact on the probability of *subjective* BR dominance being reported (see Figure 4C), but no overall impact on *objective* sensitivity to the positions of probes embedded in rivalrous inputs (see Figure 4D). Moreover, the ratings typically used by different people to describe the subjective vividness of their imagined sensations did not predict the strength of subjective (see Figure 4C) or objective (see Figure 4D) imagery priming of BR. Imagery priming in these data could not, therefore, be used to identify people who typically reported having vague or vivid visualisations. We could, however, ask a related question. Do the ratings people use to describe the vividness of their imagined experiences on *different* trials predict the degree of subjective or objective imagery priming on those trials – a within-participants effect?

To address these questions, we split participant data into low and high imagery datasets – relative to each participant’s average trial-by-trial imagery ratings. Participants who only used one imagery rating (e.g., they always reported an imagery rating of 5) were excluded from these analyses, as there were no trials with ratings that were above or below their average imagery rating. We compared the numbers of rivalrous trials resulting in imagery consistent subjective dominance reports on low vs high imagery trials. We found that participants were more likely to report imagery consistent perceptual dominance on high vs low imagery trials (t_50_ = -3.29, p = 0.002, BF_10_ = 14.9; see Figure 5).

We also compared proportions of successful position decisions regarding probes embedded in pre-imagined vs unimagined Gabors (objective imagery priming) on high vs low imagery trials. We found that objective imagery priming was greater on high vs low imagery trials (t_51_ = -2, p = 0.05, BF_10_ = 0.806; see Figure 6).

**Figure 6.**
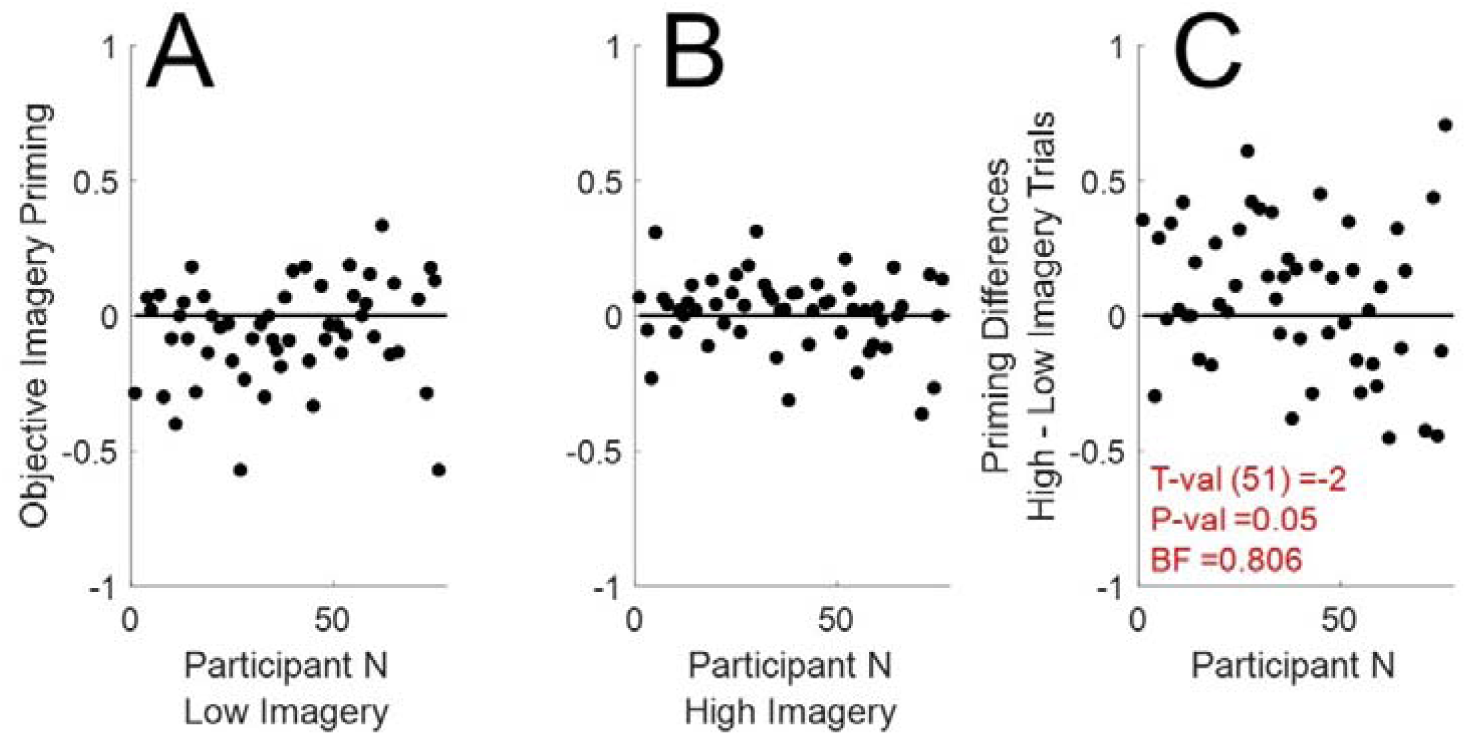
**A.** Scatterplot of imagery priming of successful probe position decisions (proportion of successful decisions regarding probes embedded in pre-imagined – unimagined Gabors, Y-axis) for different participants (X-axis) on low imagery trials. **B.** Details are as for Figure 6A, but for low imagery trials. **C.** Scatterplot of priming differences on high (see Figure 6A) vs low (see Figure 6B) imagery trials (Y-axis) for different participants (X axis).

One possible problem with our analyses is that the effects of imagery might fade over time and have little impact in the latter stages of our 10 second analysis window. We therefore conducted a second set of analyses, with a more constricted 4 second analysis window. This second set of analyses promoted a very closely matching set of results, which we present in full as supplemental material. The only modest difference was that differences in objective imagery priming on high vs low imagery trials was more robust (t_45_ = -2.37, p = 0.02, BF_10_ = 1.75; see Figure 7).

**Figure 7.**
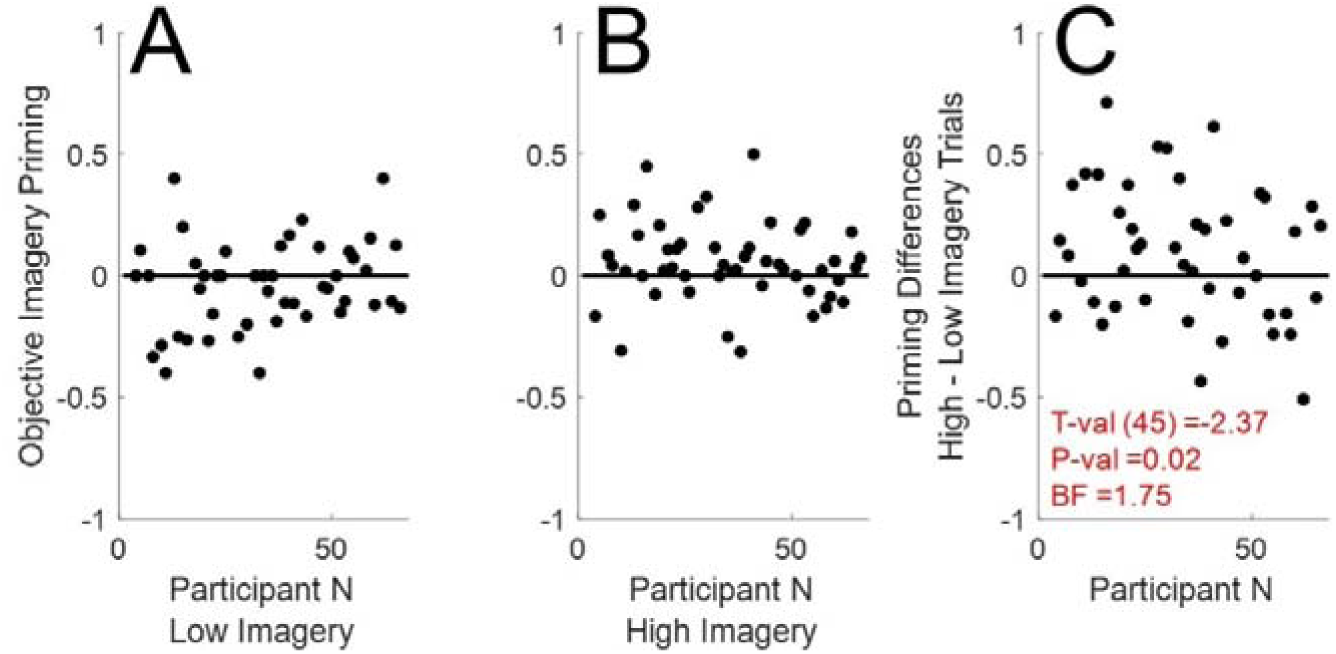
Details are as for Figure 6, but for data with time-outs recorded if perceptual dominance had not been reported after 4 seconds.

### Subjective imagery vividness ratings did not correlate with subjective BR dominance response times on non-rivalrous catch trials

Another possible correlate are the times taken to report BR dominance on non-rivalrous catch trials. While we have expressed our reservations regarding the degree to which BR can be mimicked, we nonetheless correlated the times people took to report dominance on non-rivalrous catch trials with individually averaged trial-by-trial imagery vividness ratings and VVIQ2 scores. We did this separately for non-rivalrous catch trials involving imagery consistent inputs, imagery inconsistent inputs, and the difference between these two measures. In no case was there any evidence for a significant inter-relationship (minimum P-value = 0.085, maximum BF_10_ = 0.409; see Supplemental Figure 5).

### Subjective imagery vividness ratings are more reliable than either subjective or objective imagery priming of BR

The reliability of different types of effect can be greatly informative. We therefore assessed the reliability of **1)** subjective imagery vividness ratings, **2)** imagery priming of *subjective* BR dominance reports, and **3)** imagery priming of *objective* sensitivity to probe positions, by comparing performance on randomly split halves of each participant’s trials. The ratings people used to describe their imagery were very reliable (r = 0.97, *p* < 0.001, BF_10_ > 1 million, N = 57; see Figure 8A). However, neither the extent of imagery priming of *subjective* BR dominance (r = -0.12, *p* = 0.38, BF_10_ = 0.157, N = 54; see Figure 8B) or the extent of imagery priming of *objective* sensitivity to probe positions (r = -0.007, *p* = 0.96, BF_10_ = 0.106, N = 55; see Figure 8C) were at all reliable.

**Figure 8.**
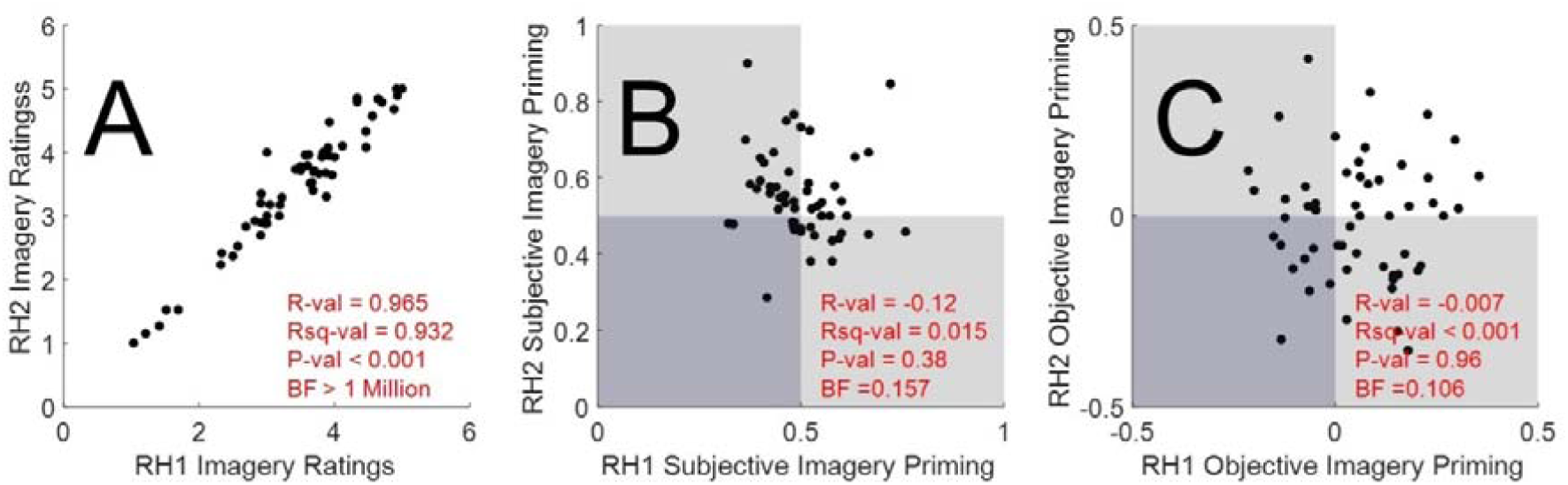
X/Y scatter plots of split-half data analyses to test for the reliability of measures. Data and analyses are depicted for split-half analyses of **A**) trial-by-trial imagery vividness ratings, **B**) Proportions of imagery consistent subjective BR dominance reports, and **C**) Imagery priming of objective sensitivity to probe positions. Datapoints within light grey shaded regions of B and C show no imagery priming for one split-half analysis. Datapoints in darker shaded regions show no imagery priming for either split-half analysis. Datapoints in unshaded regions display imagery priming for both split-half analyses.

## Discussion

Our data show that our participants were experiencing BR, as they were more sensitive to the positions of probes embedded in Gabors that were reportedly dominating perception, as opposed to Gabors that were reportedly suppressed from awareness (see Figure 4A and Supplemental Figure 3A). This replicates findings regarding the impact of binocular suppression and BR on probe sensitivity (Alais & Parker, 2006; Wales & Fox, 1970). We have also replicated an effect whereby imagery primes *subjective* BR dominance reports (see Figure 4C and Supplemental Figure 2; also see Chang et al., 2013; Pearson et al. 2008; Pearson et al., 2011).

Moreover, we have found some evidence that imagery can prime *objective* sensitivity to BR probes on a within participants basis (conceptually replicating Chang & Pearson, 2017), with people detecting a greater proportion of probes on trials on which they had reported that their visualisations had been more vivid than average for that experimental session (see Figures 6C and 7C). However, we found no robust evidence for between-participants effects, where we could predict people’s average subjective vividness ratings (or their VVIQ2 scores) from either the degree of imagery priming of *subjective* BR dominance (see Figure 4C and Supplemental Figure 2) or from the degree of imagery priming of *objective* sensitivity to probes during BR (see Figure 4D and Supplemental Figure 3B). These statements are true of our data, regardless of the time window of analysis (10 or 4 seconds).

### Imagery priming of subjective BR dominance and individual imagery vividness ratings

Our results are consistent with the majority of studies that have examined the inter-relationship between imagery priming of *subjective* BR dominance and individual imagery vividness ratings, with an absence of a robust predictive inter-relationship (e.g. Andermane et al., 2020; Bergmann et al., 2016; Dijkstra et al., 2019; Wagner & Monzel, 2024; see Table 1). Three studies have reported a robust inter-relationship. One of these had a small N (20, Pearson et al., 2011) and seems to have found an uncharacteristically large correlation relative to other studies (see Table 1). Another reported a weak interrelationship (r = 0.27) in a well powered sample (N = 105, Monzel et al., 2024) – but surprisingly did not report having to exclude *any* participants due to a bias on mock rivalry trials. A third reported a weak to moderate interrelationship (r = 0.37, N = 54, Keogh et al., 2020) and also did not report having to exclude participants due to bias on mock rivalry trials.

**Table 1.** Description of all published studies that have examined inter-relationships between imagery priming of BR and imagery vividness ratings. Type of priming refers to imagery priming of *subjective* dominance during BR (Subjective) or to imagery priming of *objective* sensitivity to BR probe positions (Objective). The Imagery questionnaire key refers to the measure of imagery vividness that imagery BR priming has been correlated with. # this correlation was calculated from data made publicly available by the authors, excluding participants with an obvious bias on mock rivalry trials (75% likely to report BR dominance on mock trials consistent or inconsistent with imagery). ## this correlation was calculated from data made publicly available by the authors.

| Study | Type of priming | N | R -value | P-value |
| --- | --- | --- | --- | --- |
| <u>Pearson et al. (2011)</u> | Subjective * | 20 |  | **<0.001** |
| <u>Andermane et al. (2020)</u> | Subjective ** | 69 | # | 0.231 |
| <u>Keogh Bergmann & Pearson</u> | Subjective *** | 54 |  | **0.003** |
| <u>Keogh Bergmann & Pearson</u> | Subjective *** | 31 |  | 0.065 |
| <u>Bergmann et al. (2016)</u> | Subjective *** | 33 |  | 0.215 |
| <u>Dijkstra et al. (2019)</u> | Subjective * | 59 |  | 0.470 |
| <u>Wagner & Monsel (2023)</u> | Subjective **** | 66 | ## | 0.118 |
| <u>Monzel et al. (2024)</u> | Subjective **** | 105 |  | **0.005** |
| **Bouyer et al.** | Subjective *** / * | 56 |  | 0.670 |
|  | Objective *** / * | 57 |  | 0.890 |
| **Imagery Questionnaires: \* VVIQ2 \*\*Sussex Cognitive Styles Questionnaire, Imagery Ability subscale \*\*\* Trial-by-trial Imagery Vividness Ratings \*\*\*\* VVIQ** |  |  |  |  |

In sum, we believe the balance of existing evidence suggests that imagery priming of *subjective* BR dominance remains open to question. The average effect reported across all studies suggests a weak inter-relationship (r = 0.271) might exist which would have little utility as a tool to identify if a particular person might have a weak or a non-existent capacity to visualise (as indexed by imagery vividness ratings).

Moreover, we stress that by nature this evidence is *subjective*, so it is not clear if this evidence is a conceptual advance on simply asking people directly about the vividness of their visualisations.

To be clear, a large number of studies have shown that imagery *can* prime subjective BR dominance reports (e.g. Andermane et al., 2020; Chang et al., 2013; Dijkstra et al., 2019; Keogh & Pearson, 2011, 2014, 2017, 2018; Keogh et al., 2020; Maróthi & Kéri, 2018; Pace et al., 2023; Rademaker & Pearson, 2012; Shine et al., 2015), and we have replicated that group level effect here (see Figure 4C and Supplemental Figure 2). A number of studies have also shown group level differences between the extent of imagery priming of BR (e.g. Shine et al., 2015).

Table 1 and our discussion above do not question these effects. The issue is rather if the extent of imagery priming of *subjective* BR dominance reports can be used to reliably predict individual differences in imagery vividness ratings. We do not believe there is good evidence that it can.

### Imagery Priming of Objective sensitivity to probes during BR

Ours is the first study to attempt to correlate imagery priming of *objective* sensitivity to probes during BR with individual differences in imagery vividness ratings. We have found no evidence for a robust inter-relationship between these two measures – which would have been a between-participants effect.

We did find some evidence that people were more sensitive to the positions of probes embedded in imagery consistent Gabors on trials when they had rated their imagery as more vivid than average (for them during that experimental session, see Figure 7). So, there is some evidence for a within-participants effect, whereby people are more sensitive to imagery primed probes when they feel that their visualisations had been more vivid than usual – conceptually replicating Chang & Pearson (2017).

### The VVIQ2

Currently the most popular subjective metric of the vividness of imagery is the VVIQ2 (Marks, 1995). But there is a growing consensus that we need a better means of assessing individual differences in the vividness of our imagined visual sensations (Blomkvist & Marks, 2023; Schwarzkopf, 2024). It has been hoped that imagery priming of BR could provide a replacement metric. To date, we would argue that variants of the BR protocol have fallen short of that goal. On average across published studies (see Table 1) there is only evidence for a weak correlation between imagery priming of subjective BR dominance and VVIQ2 scores (r ∼0.271). Such a weak association would have little utility as a tool to identify people who are likely to have a weak or an absent capacity to visualise (Zeman et al., 2015).

Moreover, we emphasise that imagery priming of BR dominance is a *subjective* effect, and consequently that it is not clear that this is a conceptual advance on simply asking people about the vividness of their visualisations.

Ours is the first study to attempt to correlate imagery priming of *objective* sensitivity to probe positions during BR with individual differences in the vividness of imagery.

We did not find any evidence for a robust correlation between these measures (see Figure 4D, and Supplemental Figure 3B).

There are other candidates to displace subjective questionnaires as a metric of the vividness of imagined visual experiences. For instance, Kay et al. (2022) have suggested that pupil dilation to imagined bright or dark images could be used as a physiological index of the intensity of imagined visual experiences. We plan to examine this effect in a future study.

### Which is a worse measure of the vividness of imagery, Imagery Priming of BR or VVIQ2 scores?

One question that can reasonably be asked is which was a worse predictor of individual differences in the subjective vividness of imagery. Was it imagery priming of subjective BR dominance, or was it the subjective ratings different people use to describe their imagery? On the basis of some of our data, and similar datasets (e.g. Andermane et al., 2020; Bergmann et al., 2016; Dijkstra et al., 2019; Wagner & Monzel, 2024), all we can claim is that these measures are not reliable predictors of one another.

We can, however, consider the internal reliability of these measures, as we have done in Figure 8. This analysis reveals that the ratings people used to describe the vividness of their visualisations were very reliable (see Figure 8A), whereas neither the extent of imagery priming of *subjective* BR dominance reports (see Figure 8B) or the extent of imagery priming of *objective* sensitivity to probe positions (see Figure 8C) were at all reliable. It is admittedly difficult to interpret these data, as we do not know precisely how reliable different people’s experiences of imagery are. Some evidence suggests people’s experiences of visual imagery can fluctuate over time within a trial, and are variable across trials (see Sulfaro et al., 2024). But how variable is imagery across trials? If imagery is reasonably consistent across trials, our data would suggest that trial-by-trial vividness ratings are likely the best of our candidate measures of the vividness of visualisations. If imagery is highly variable across trials, then it is possible that one or both of the BR-based measures is a better measure.

### Does imagery priming of subjective BR dominance measure of the vividness of imagery, or a related cognitive process?

Another interpretive ambiguity is if the act of imagery has directly primed subjective dominance during BR (see Figure 4 and Supplemental Figure 2; also see Andermane et al., 2020; Chang et al., 2013; Dijkstra et al., 2019; Keogh & Pearson, 2011, 2014, 2017, 2018; Keogh et al., 2020; Maróthi & Kéri, 2018; Pace et al., 2023; Rademaker & Pearson, 2012; Shine et al., 2015), or if subjective BR dominance reports have been biased by a cognitive operation that is inter-linked with imagery – such as awareness of the semantic descriptions of imagined content. Enhanced performance on probe position decisions (see Figures 7 – 8; Chang & Pearson, 2017) is less likely to be explicable as a form of semantic priming of visual content, as the information that is needed for a correct probe position decision is unrelated to the semantics of the imagined content.

### Study limitations

One feature that may have weakened the impact of imagery in our study was the balanced number of catch trials. These meant that there was no rivalry on 50% of trials. Because of this, participants might have learnt that their imagination had little impact on their visual experiences. This could have resulted in a learnt helplessness situation (Teodorescu & Erev, 2014), with people giving up on trying to control rivalry outcomes with their imagination. While we acknowledge this possibility, our results show clear evidence of imagery priming of *subjective* BR dominance (see Figure 4 and Supplemental Figure 2), and some evidence for imagery priming of *objective* probe sensitivity (see Figures 7 – 8). So, our evidence suggests that people were engaging in imagery, and that this was modulating their visual experiences. In our data, however, these effects were not associated with individual differences in imagery vividness ratings (see Figure 4C-D and Supplemental Figures 2 and 3B).

Another potential limitation is that we did not conduct a preliminary task to individually calibrate BR inputs to correct for eye dominance (e.g. Pearson et al., 2008). This term can refer to an individualized tendency to report that inputs shown to a particular eye have dominated perception during BR (Porac & Coren, 1976), independent of any additional bias resulting from other factors – such as pre-imagining a rivalrous input. The effects of eye dominance can reportedly be counteracted by adjusting the relative contrasts of rivalrous inputs, to arrive at a point of equilibrium (e.g. Pearson et al., 2008). We are not aware of any data that attests to the reliability of this procedure, but a potentially related BR based measure of eye dominance has been shown to be unreliable (Min et al., 2022). Nonetheless, if the effects of eye dominance could reliably be countermanded, this might provide an optimal basis from which to study the impact of additional biasing factors – such as imagery. However, we do not believe the omission of an eye dominance calibration task has been a serious limitation for our study.

If eye dominance had been the cause of our failure to detect robust inter-relationships between imagery priming of BR, we would expect to see a trend toward greater evidence for inter-relationships as we impose tighter exclusion criteria for eye dominance (relative to the 90% criterion we have used in our main analyses).

However, in our data we see no such trend. For criteria ranging from 60 to 90% (in 10% steps), bayes factors relating to *both* imagery priming of *subjective* BR dominance and to objective sensitivity to BR probes all provide moderate levels of evidence in favour of null hypotheses, that imagery priming of BR is unrelated to measures of the subjective vividness of imagery (see Supplemental Table 1).

### Conclusions

Finding a reliable objective metric of the vividness of visualisations would not just assuage a conceptual curiosity. It could have practical implications. Visual imagery is used among the world’s most popular psychological interventions to benefit mental health (e.g., Odou & Vella-Brodrick, 2013) and performance (Jones & Stuth, 1997). We argue that we do not currently have a well validated objective predictor of the subjective vividness of people’s visualisations, so we cannot be sure that all people have equal potential to benefit from such interventions.

We have examined the impact of pre-imagining rivalrous inputs on *subjective* BR dominance reports, and more importantly for the first time on an *objective* measure of sensitivity to probes embedded in BR inputs. We have found some evidence that pre-imagining a rivalrous input can prime both outcomes at a group level basis.

However, in our data neither the degree of imagery priming of *subjective* BR dominance, or the degree of imagery priming of *objective* sensitivity to probes embedded in BR inputs, was able to predict individual differences in imagery vividness ratings. We argue that further work is needed to identify a reliable objective approach to predicting individual differences in the subjective vividness of imagery.

### Ethics

Ethical approval was obtained from the University of Queensland’s (UQ) Ethics Committee, and the experiment was performed in accordance with UQ guidelines and regulations for research involving human participants. Each participant provided informed consent to participate in the study and were made aware that they could withdraw from the study at any moment without prejudice or penalty.

### Declaration of Competing Interest

The authors declare that they have no known competing financial interests or personal relationships that could have appeared to influence the work reported in this paper.

### Data availability

Data and analysis scripts are publicly available.

## Notes

https://search.library.uq.edu.au/permalink/f/18av8c1/61UQ_eSpace00effb3

### Competing Interest Statement

The authors have declared no competing interest.

## References

Andermane, N., Bosten, J.M., Seth, A. & Ward, J. (2020). Individual differences in the tendency to see the expected. Consciousness and Cognition, 85, 102989 10.1016/j.concog.2020.102989

Anderson, J. D., Bechtoldt, H. P., & Dunlap, G. L. (1978). Binocular integration in line rivalry. Bulletin of the Psychonomic Society, 11, 399–402.

Arnold, D.H. (2011). Why is binocular rivalry uncommon? Frontiers in Human Neuroscience 5: 116 doi: 10.3389/fnhum.2011.00116

Arnold, D. H., Andresen, I., Anderson, N., & Saurels, B. W. (2023). Commonalities between the Berger Rhythm and spectra differences driven by cross-modal attention and imagination. Conscious Cogn, 107, 103436. 10.1016/j.concog.2022.103436

Arnold, D. H., Wegener, S. V., Brown, F., & Mattingley, J. B. (2012). Precision of synesthetic color matching resembles that for recollected colors rather than physical colors. Journal of Experimental Psychology. . Human Perception and Performance, 38(5), 1078–1084. 10.1037/a0028129

Barron, H. C., Auksztulewicz, R., & Friston, K. (2020). Prediction and memory: A predictive coding account. Prog Neurobiol, 192, 101821. 10.1016/j.pneurobio.2020.101821

Barron, H. C., Dolan, R. J., & Behrens, T. E. (2013). Online evaluation of novel choices by simultaneous representation of multiple memories. Nat Neurosci, 16(10), 1492–1498. 10.1038/nn.3515

Bergmann, J., Genç, E., Kohler, A., Singer, W., & Pearson, J. (2016). Smaller Primary Visual Cortex Is Associated with Stronger, but Less Precise Mental Imagery. Cerebral Cortex, 26(9), 3838–3850. 10.1093/cercor/bhv186

Blomkvist, A., & Marks, D. F. (2023). Defining and ’diagnosing’ aphantasia: Condition or individual difference? Cortex, 169, 220–234. 10.1016/j.cortex.2023.09.004

Brascamp, J. W., Klink, P. C., & Levelt, W. J. M. (2015). The ’laws’ of binocular rivalry: 50 years of Levelt’s propositions. Vision Research, 109, 20–37. 10.1016/j.visres.2015.02.019

Brown, R. J. & Norcia, A. M. (1997). A method for investigating binocular rivalry in real-time with steady-state VEP. Vision Research 37, 1401–2408.

Chang, S., & Pearson, J. (2018). The functional effects of prior motion imagery and motion perception. Cortex, 105, 83–96. 10.1016/j.cortex.2017.08.036

Chang, S., Lewis, D.E. & Pearson J. (2013). The functional effects of color perception and color imagery. Journal of Vision 13(10): 4. 10.1167/13.10.4

Cui, X., Jeter, C. B., Yang, D., Montague, P. R., & Eagleman, D. M., 47(4), . . (2007). Vividness of mental imagery: Individual variability can be measured objectively. . Vision Research (Oxford), 47(4), 474–478. 10.1016/j.visres.2006.11.013

Dijkstra, N., Bosch, S. E., & van Gerven, M. A. J. (2017). Vividness of Visual Imagery Depends on the Neural Overlap with Perception in Visual Areas. Journal of Neuroscience, 37(5), 1367–1373. 10.1523/Jneurosci.3022-16.2016

Dijkstra, N., Hinne, M., Bosch, S.E. & van Gerven, M.A.J. (2019). Between-subject variability in the influence of mental imagery on conscious perception. Scientific Reports 9, 15658. 10.1038/s41598-019-52072-1

Dijkstra, N., Kok, P., & Fleming, S. M. (2022). Perceptual reality monitoring: Neural mechanisms dissociating imagination from reality. Neuroscience and Biobehavioral Reviews, 135. ARTN 104557 10.1016/j.neubiorev.2022.104557

Gallagher, R., Suddendorf, T. & Arnold, D.H. (2019). Confidence as a diagnostic tool for perceptual aftereffects. Scientific Reports 9, 7124.

Galton, F. (1880). Statistics of Mental Imagery. Mind, 5(19), 19.

Green, D.M. & Swets, J.A. (1966). Signal detection theory and psychophysics Wiley, New York.

Grzanka, P. R., & Moradi, B. (2021). The qualitative imagination in counseling psychology: Enhancing methodological rigor across methods. J Couns Psychol, 68(3), 247–258. 10.1037/cou0000560

Haynes, J. & Rees, G. (2005). Predicting the stream of consciousness from activity in human visual cortex. Current Biology, 15, 1301–1307.

Hirsch, C. R., & Holmes, E. A. (2007). Mental imagery in anxiety disorders. Psychiatry (Abingdon, England), 6(4), 161–165. 10.1016/j.mppsy.2007.01.005

Holmes, E. A., & Mathews, A. (2010). Mental imagery in emotion and emotional disorders. Clinical Psychology Review, 30(3), 349–362. 10.1016/j.cpr.2010.01.001

Jones, L., & Stuth, G. (1997). The uses of mental imagery in athletics: An overview. Applied & Preventive Psychology, 6(2), 101–115. 10.1016/S0962-1849(05)80016-2

Jozwik, K. M., O’Keeffe, J., Storrs, K. R., Guo, W. X., Golan, T., & Kriegeskorte, N. (2022). Face dissimilarity judgments are predicted by representational distance in morphable and image-computable models. Proceedings of the National Academy of Sciences of the United States of America, 119(27). ARTN e2115047119 10.1073/pnas.2115047119

Kay, L., Keogh, R., Andrillon, T., Pearson, J., & Serences, J. T. (2022). The pupillary light response as a physiological index of aphantasia, sensory and phenomenological imagery strength. Elife, 11. ARTN e72484 10.7554/eLife.72484

Keogh R, Bergmann J, Pearson J. (2020). Cortical excitability controls the strength of mental imagery. Elife. 9, e50232. 10.7554/eLife.50232

Keogh, R. & Pearson, J. (2011). Mental imagery and visual working memory. PLoS One. 6, e29221. 10.1371/journal.pone.0029221

Keogh, R. & Pearson, J. (2014). The sensory strength of voluntary visual imagery predicts visual working memory capacity. Journal of Vision 14(12), 7. 10.1167/14.12.7

Keogh, R. & Pearson, J. (2017). The perceptual and phenomenal capacity of mental imagery. Cognition 162, 124–132. 10.1016/j.cognition.2017.02.004

Keogh, R. & Pearson, J. (2018). The blind mind: No sensory visual imagery in aphantasia. Cortex 105, 53–60. 10.1016/j.cortex.2017.10.012

Kwok, E. L., Leys, G., Koenig-Robert, R., & Pearson, J. (2019). Measuring Thought-Control Failure: Sensory Mechanisms and Individual Differences. Psychological Science, 30(6), 811–821. 10.1177/0956797619837204

Marks, D.F. (1995). New Directions for Imagery Research. Journal of Mental Imagery 19, 153–167.

Min, S.H., Gong, L., Baldwin, A.S., Reynaud, A., He, Z., Zhou, J. & Hess, R.F. (2021). Some psychophysical tasks measure ocular dominance plasticity more reliably than others. Journal of Vision, 21(8), 20. 10.1167/jov.21.8.20

Morina, N., Leibold, E., & Ehring, T. (2013). Vividness of general mental imagery is associated with the occurrence of intrusive memories. J Behav Ther Exp Psychiatry, 44(2), 221–226. 10.1016/j.jbtep.2012.11.004

Maróthi, R. & Kéri, S. (2018). Enhanced mental imagery and intact perceptual organization in schizotypal personality disorder. Psychiatry Research, 259, 433–438. 10.1016/j.psychres.2017.11.015

Odou, N. & Vella-Brodrick, D. A. (2013). The efficacy of positive psychology interventions to increase well-being and the role of mental imagery ability. Social Indicators Research, 110(1), 111–129. 10.1007/s11205-011-9919-1

Pace, T., Koenig-Robert, R. & Pearson, J. (2023). Different Mechanisms for Supporting Mental Imagery and Perceptual Representations: Modulation Versus Excitation. Psychological Science 34, 1229–1243. 10.1177/09567976231198435

Pearson, J. (2019). The human imagination: the cognitive neuroscience of visual mental imagery. Nature Reviews Neuroscience 20(10), 624–634. 10.1038/s41583-019-0202-9

Pearson, J. Clifford, C.W.G. & Tong, F. (2008). The Functional Impact of Mental Imagery on Conscious Perception. Current Biology, 18(13), 982–986. 10.1016/j.cub.2008.05.048

Pearson, J., Rademaker, R.L. & Tong, F. (2011). Evaluating the Mind’s Eye. Psychological Science 22, 1535–1542.

Porac, C., & Coren, S. (1976). The dominant eye. Psychological Bulletin 83, 880 – 897. 10.1037/0033-2909.83.5.880

Rademaker, R.L. & Pearson, J. (2012). Training visual imagery: improvements of metacognition, but not imagery strength. Frontiers in Psychology 3, 224 10.3389/fpsyg.2012.00224

Reddan, M. C., Wager, T. D., & Schiller, D. (2018). Attenuating Neural Threat Expression with Imagination. Neuron, 100(4), 994-+. 10.1016/j.neuron.2018.10.047

Roseboom, W. & Arnold, D.H. (2011). Twice upon a time: Multiple, concurrent, temporal recalibrations of audio-visual speech. Psychological Science 22, 872–877.

Sulfaro, A.A., Robinson, A.K. and Carlson, T.A. (2024). Properties of imagined experience across visual, auditory, and other sensory modalities, Consciousness and Cognition 117, 103598. DOI:10.1016/j.concog.2023.103598.

Shine, J.M., Keogh, R., O’Callaghan, C., Muller, A.J., Lewis, S.J. & Pearson J. (2015). Imagine that: elevated sensory strength of mental imagery in individuals with Parkinson’s disease and visual hallucinations. Proceedings of the Royal Society of London B, 282, 20142047, 1798 10.1098/rspb.2014.2047

Schacter, D. L., Addis, D. R., Hassabis, D., Martin, V. C., Spreng, R. N., & Szpunar, K. K. (2012). The Future of Memory: Remembering, Imagining, and the Brain. Neuron, 76(4), 677–694. 10.1016/j.neuron.2012.11.001

Schwarzkopf, D. S. (2024). What is the true range of mental imagery? [Viewpoint]. Cortex, 170, 21–25. 10.1016/j.cortex.2023.09.013

Storrs, K. R. (2015). Are high-level aftereffects perceptual? Frontiers in psychology, 6. 10.3389/fpsyg.2015.00157

Teodorescu, K., & Erev, I. (2014). Learned Helplessness and Learned Prevalence: Exploring the Causal Relations Among Perceived Controllability, Reward Prevalence, and Exploration. Psychological Science, 25(10), 1861–1869. 10.1177/0956797614543022

Veraksa, A. N., & Gorovaya, A. E. (2011). Effect of Imagination on Sport Achievements of Novice Soccer Players. Psychology in Russia-State of the Art, 4, 495–504.<GO to ISI>://WOS:000414894000033

Wendt, G. & Faul, F. (2022). Binocular luster – A review. Vision Research, 194, 108008. 10.1016/j.visres.2022.108008

Yarrow, K., Jahn, N., Durant, S. & Arnold, D.H. (2011). Shifts of criteria or neural timing? The assumptions underlying timing perception studies. Consciousness & Cognition, 20, 1518–1531.

Zeman, A., Dewar, M., & Della Sala, S. (2015). Lives without imagery - Congenital aphantasia. Cortex, 73, 378–380. 10.1016/j.cortex.2015.05.019

Zeman, A. (2024). Aphantasia and hyperphantasia: exploring imagery vividness extremes. Trends in Cognitive Sciences, 28(5), 467–480. 10.1016/j.tics.2024.02.007

