## Supplementary Material for "Imagery priming of binocular rivalry is not a reliable metric of individual differences in the subjective vividness of visualisations"

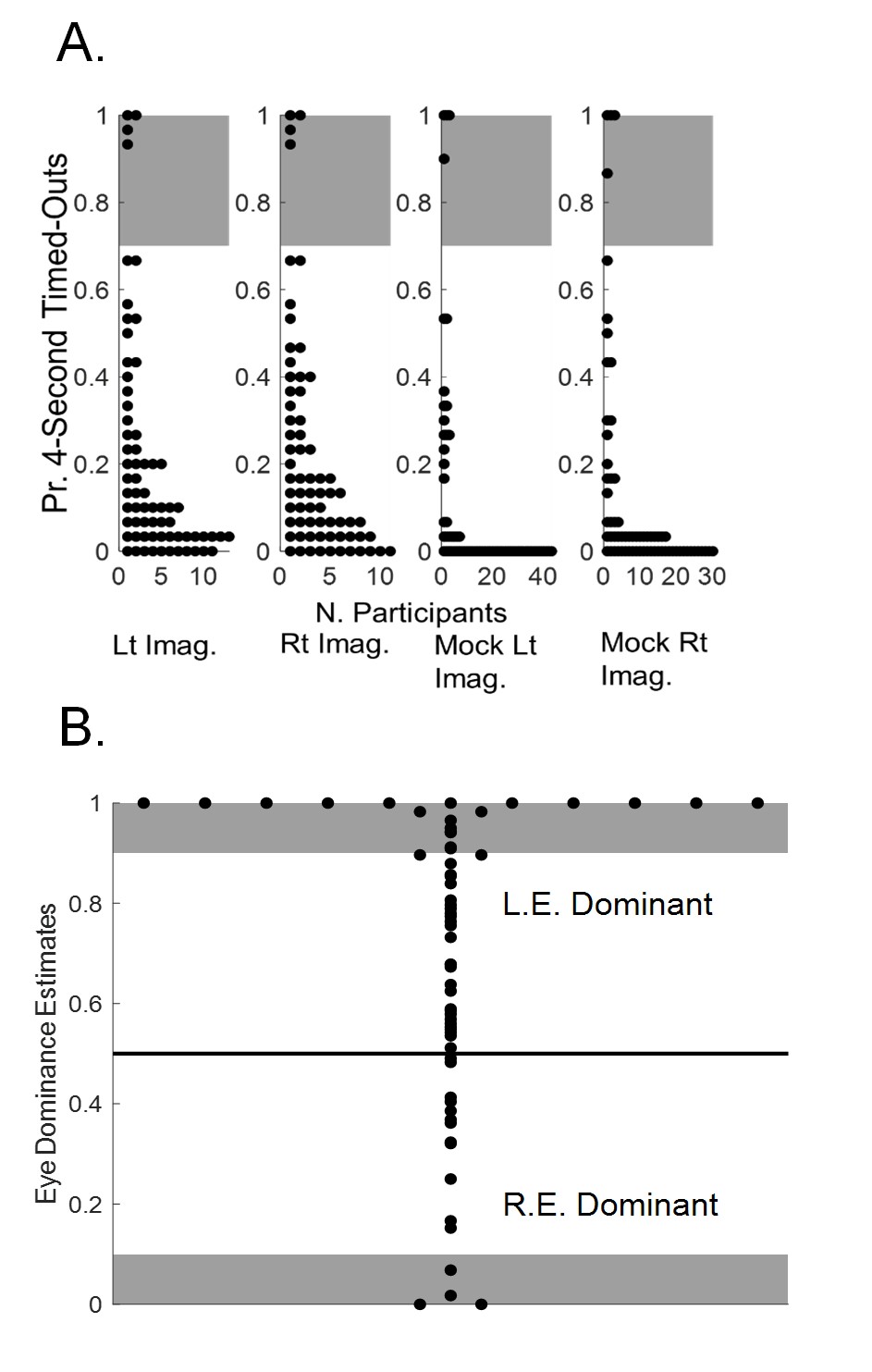

**Supplemental Figure 1.** Details are as for paper Figure 3, with the exception that time-outs were recorded if perceptual dominance had not been reported after 4 seconds.

For our second set of analyses, participants who timed-out (who did not report dominance) after 4 seconds on 70% or more of rivalrous presentations were excluded from further analyses (see Supplemental Figure 1A). We then re-calculated eye dominance estimates, as many trials that had been included in our original eye dominance calculations had been excluded from this second set of analyses. So again, we excluded participants from further analyses if they had reported dominance of the Gabors shown to one eye (left or right) on 90% or more of rivalrous presentations (see Supplemental Figure 1B). These initial exclusions resulted in a potential dataset of 52 participants. As per our initial set of analyses, we also excluded datapoints from any analyses that were +-3 standard deviations from the conditional mean, and we only included participant data in a conditional analysis if at least five viable trials had informed that datapoint. The precise number of participants informing each analysis can again be inferred from the reported degrees of freedom for each test.

People were again more likely to report perceptual dominance of pre-imagined Gabors (t_50_ = 2.56, p = 0.01, BF_10_ = 2.63; see Supplemental Figure 2). However, we still did not find any evidence of a relationship between imagery priming of perceptual dominance and either the average ratings people used to describe the subjective intensity of their pre-imagined Gabors (r = 0.05, *p* = 0.71, N = 51; see Supplemental Figure 2) or their VVIQ2 scores (r = -0.04, p = 0.76).

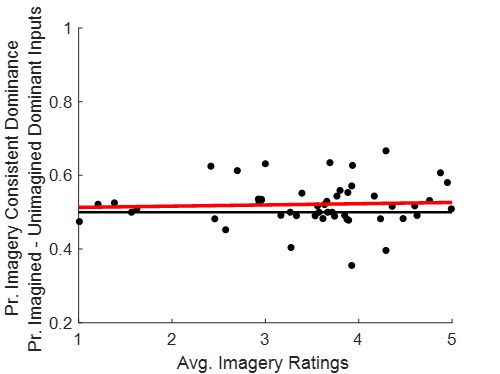

**Supplemental Figure 2.** Details are as for paper Figure 4, but for data with time-outs recorded if perceptual dominance had not been reported after 4 seconds.

We also again found that that people made more successful probe position decisions when these were embedded in Gabors that were reportedly currently dominating perception, as opposed to Gabors that were reportedly currently suppressed from awareness (paired t_49_ = 13.7, p < 0.001; see Supplemental Figure 3A). A Bayes Factor analysis revealed extreme evidence for the alternative hypothesis in this case, that people would make more successful probe position decisions about probes embedded in reportedly currently dominating perception (BF_10_ > 1 million). There was, however, no evidence that these effects were related to the average ratings different people used to describe the subjective intensity of their pre-imagined Gabors (r = -0.01, *p* = 0.99, N = 50; see Figure 4). There was, however, a negative relationship between perceptual dominance priming of successful probe position decisions and individual VVIQ2 scores (r = -0.29, p = 0.04).

People did not make proportionally more successful decisions about the positions of probes embedded in pre-imagined as opposed to unimagined Gabors (t_50_ = 1.45, p = 0.2, BF_10_ = 0.36; see Supplemental Figure 3B), and there was no evidence for a relationship between imagery priming of successful probe position decisions and either the average imagery ratings people used to describe the vividness of their pre-imagined Gabors (r = 0.11, p = 0.45; see Supplemental Figure 3B) or their VVIQ2 scores (r = -0.04, p = 0.76).

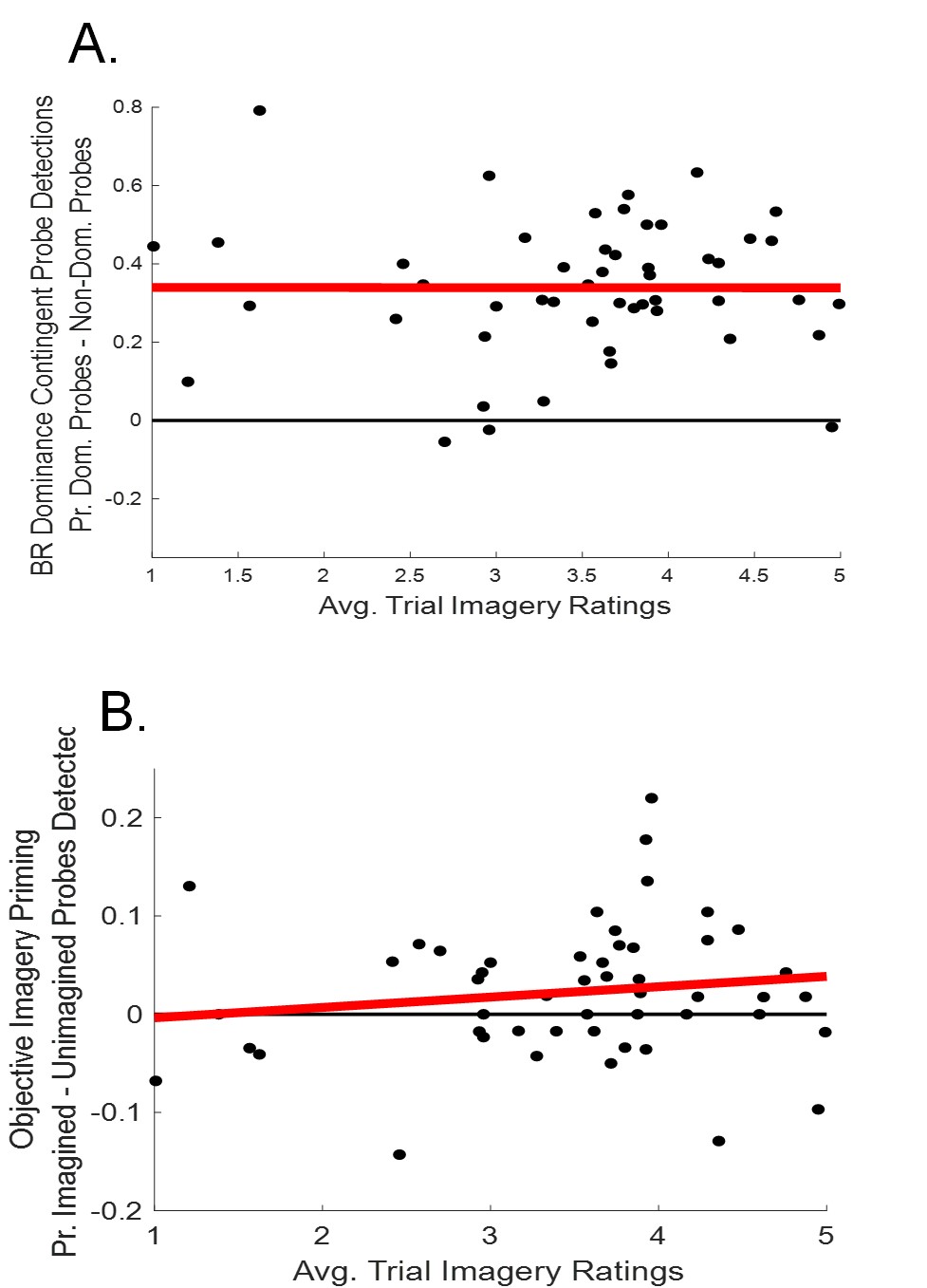

**Supplemental Figure 3.** Details are as for paper Figure 5, but for data with time-outs recorded if perceptual dominance had not been reported after 4 seconds.

When we split these data by the average rating each participant used to describe the vividness of their experiences of pre-imagined Gabors, we again found that participants were more likely to report imagery consistent perceptual dominance on high vs low imagery trials (t_44_ = -2.73, p = 0.009, BF_10_ = 3.72; see Supplemental Figure 4).

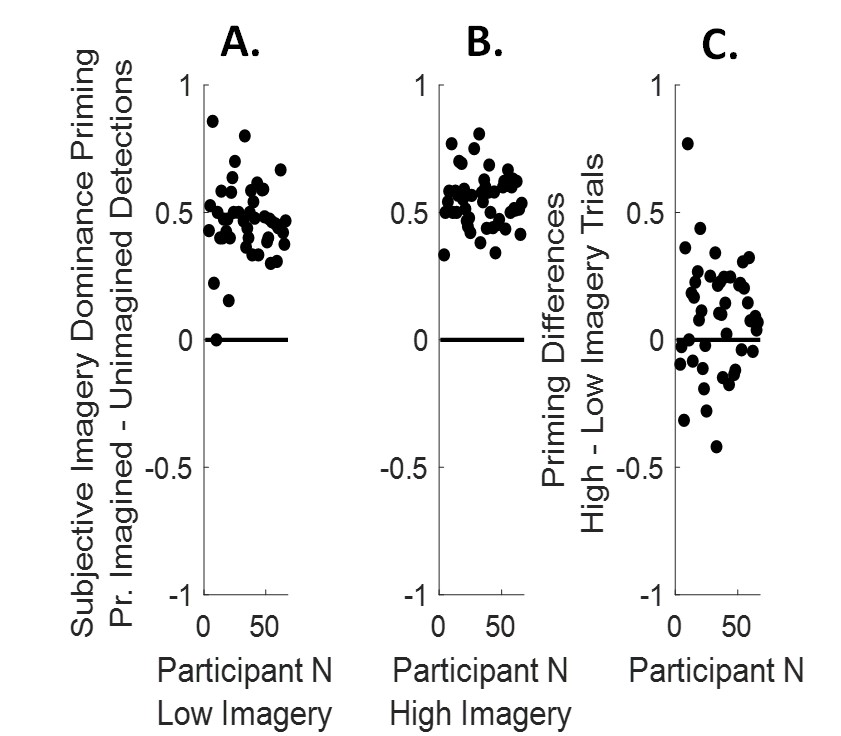

**Supplemental Figure 4.** Details are as for paper Figure 6, but for data with time-outs recorded if perceptual dominance had not been reported after 4 seconds.
